# PANAMA enabled high sensitivity dual nanoflow LC/MS metabolomics and proteomics analysis

**DOI:** 10.1101/2023.11.03.565531

**Authors:** Weiwei Lin, Fatemeh Mousavi, Benjamin C. Blum, Christian F. Heckendorf, Noah Lampl, Ryan Heckman, Hongbo Guo, Mark McComb, Andrew Emili

## Abstract

High sensitivity nanoflow liquid chromatography (nLC) is seldom employed in untargeted metabolomics because current sample preparation techniques are inefficient to prevent nanocapillary column performance degradation. Here, we describe an nLC-based tandem mass spectrometry workflow that enables seamless joint analysis and integration of metabolomics (including lipidomics) and proteomics from the same samples without instrument duplication. This workflow is based on robust solid phase micro-extraction step for routine sample clean-up and bioactive molecule enrichment. Our method, termed PANAMA, improves compound resolution and detection sensitivity without compromising depth of coverage as compared with existing widely used analytical procedures. Notably, PANAMA can be applied to a broad array of specimens including biofluids, cell line and tissue samples. It generates high quality, information rich metabolite-protein datasets while bypassing the need for specialized instrumentation.

**Motivation:** The ability to routinely, sensitively and reproducibly analyze both cellular proteins and metabolite mixtures from the same biospecimens can enhance the discovery of biomolecules associated with basic biochemical processes and pathobiological states. Yet existing mass spectrometry-based profiling methods rely on specialized protocols and duplicated instrumentation platforms, resulting in increased time, sample consumption and costs. We sought to generate an effective platform for both metabolomic and proteomic studies on the same samples by enabling nanoflow liquid chromatography for small molecules. The resulting approach was extensively optimized and benchmarked to provide in depth molecular coverage, along with improved chromatographic separations, sensitivity and reliability as compared to existing methods. The cost benefit ratio of PANAMA is substantial because the platform bypasses the need for specialized instrumentation stemming from incompatible procedures.

## Introduction

Proteomics and metabolomics provide complementary insights into the biochemical pathways that are altered by pathophysiological conditions or disease stages^1^. Specialized high- performance liquid chromatography-mass spectrometry (LC-MS) are usually used for such analyses as the physicochemical properties of metabolites and lipids are distinct from those of proteins and peptides^2,3^. Most notably, whereas nanoflow liquid chromatography-mass spectrometry (nLC-MS) is widely applied in proteomics due to its high sensitivity^4^, it is seldom employed for metabolomics, including lipidomics. While several attempts have been reported relying on either a syringe pump or a static nanoESI source^5,6^, nLC separation is still rarely deployed for untargeted small molecule surveys.

A key bottleneck in current sample preparation techniques is the inefficient removal of matrix components (e.g. salts, phospholipids, intact proteins) that tend to clog nanocapillary columns, leading to rapid degradation in LC-MS performance (e.g. nanospray instability, ion suppression)^7^. Current analyte sample preparation procedures, such as organic solvent precipitation or liquid-liquid extraction (LLE), are typically optimized for high (microflow) downstream analyses (mLC)^8^. Hence, while nLC electrospray ionization can potentially enhance the resolution and detection of low abundance analytes, efficient sample clean-up is crucial to avoid nanocolumn obstruction. In essence, existing methodologies warrant separate preparation and analysis instrumentation for metabolites, proteins and lipids, while consuming a higher quantity of biomaterial. This becomes a critical limitation for translation research since human samples are frequently available in limiting amounts (e.g. tissue biopsies).

Building upon well-established proteomic and modern solid phase micro-extraction (SPME) workflows, we have devised, optimized and benchmarked a unified <u>P</u>roteomic <u>a</u>nd <u>Na</u>noflow <u>M</u>etabolomic <u>A</u>nalysis workflow (PANAMA) that minimizes capillary blockage allowing for the reproducible detection and measurement of a wide range of metabolites, lipids and proteins of broad biological interest from complex samples, such as blood, on single nLC/MS platform. We show that this integrated pipeline provides comparable or even improved chromatographic separation and quantitative coverage of diverse biomolecules compared with that obtained using standard independent metabolomic workflows^9,10^. The PANAMA platform can be successfully employed on a broad array of specimens including biofluids, cell lines and tissues, as described in our other recent publications^11,12^.

## Results

### SPME-assisted sample clean-up enables dual metabolomic-proteomic nanoflow LC/MS

To facilitate joint protein, lipid and metabolite analysis using a single standardized instrumentation platform, we adapted and optimized an SPME sample preparation protocol^13^ suitable for routine untargeted metabolomics analysis by nLC-MS that we found avoids nanocolumn obstruction. As outlined schematically in **Fig. 1**, complex biospecimens (biofluids, animal tissues, cultured cells) are first subject to liquid-liquid extraction (LLE) using cold ACN/MeOH/H_2_O (4/4/2, v/v) to precipitate proteins while bulk extracting cellular metabolites. (alternatively, cold H_2_O:MeOH:CHCl_3_ (50/45/5,v/v) can be used). Then, after sample concentration (SpeedVac), the supernatant (metabolites) is processed using SPME to eliminate confounding matrix contaminants (**Fig.1a**). Throughput can be enhanced using a microplate compatible 96-blade autosampler device (**Fig. 1b-iv,** see also **Supplementary Video 1**). Broad capture of both polar and non-polar compounds (-7<logP<15) is achieved by inserting thin metallic blades coated with a 1:1 (w/w) hydrophilic-lipophilic sorbent mixture consisting of polystyrene-divinylbenzene and a weak anion exchanger (PS-DVB-WAX, CHROMABOND, Germany) and Polypropylene (HLB, Waters, USA) ^13,14^ (**Fig. 1b-i/ii**; see also **Supplementary Protocol 1**). Salts, intact proteins are removed by solvent precipitation are removed by extensive washing of the blades (**Fig. 1b-iii**).

**Figure 1.**
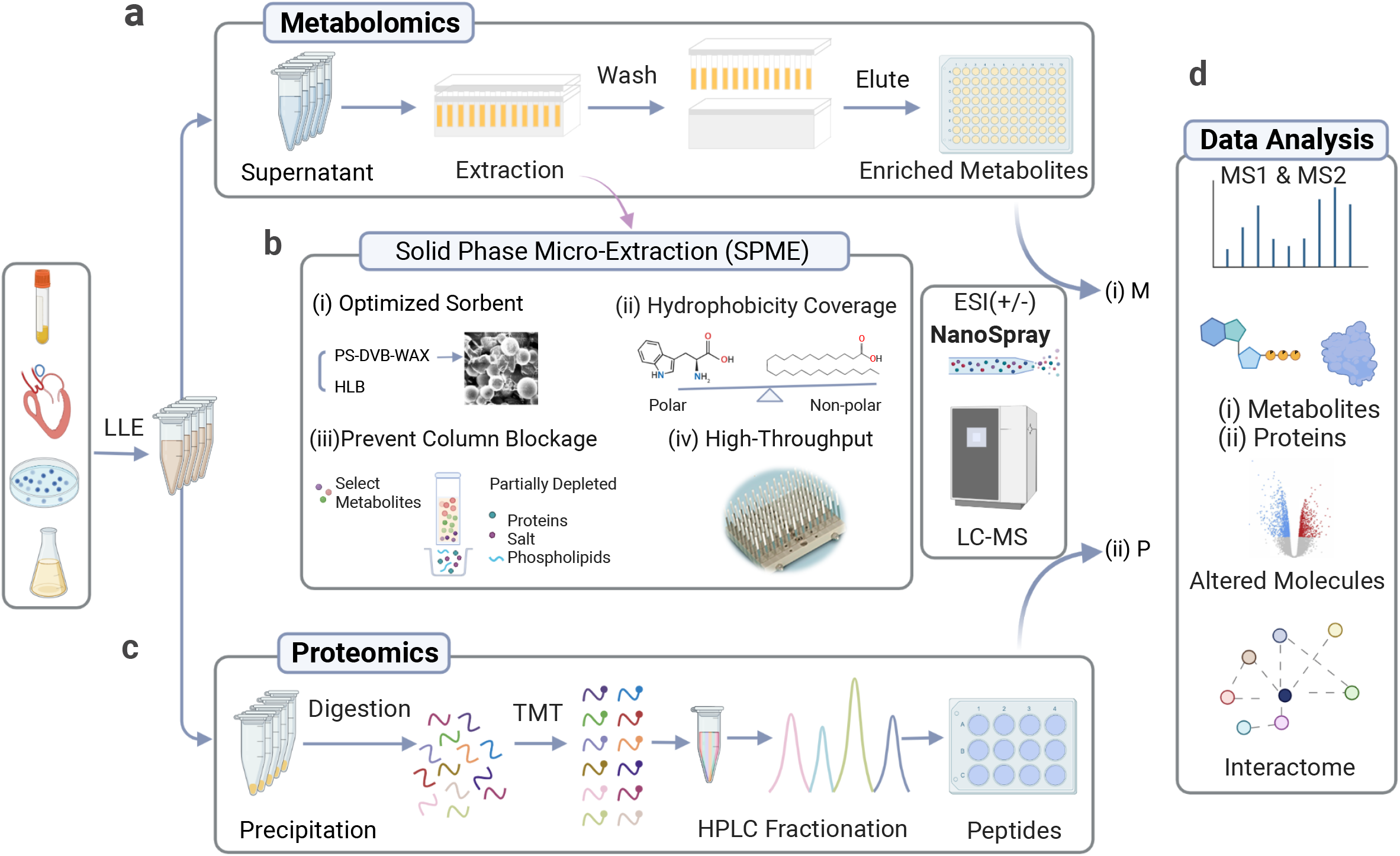
PANAMA enables joint metabolite-lipid-protein profiling from single samples using a unified SPME-nLC/MS workflow Schematic illustrating key steps in PANAMA workflow: **a**, After organic solvent extraction, metabolites are cleaned up and concentrated by SPME prior to nanoflow LC/MS analysis with positive/negative ion mode switching. **b**, Diverse metabolites are captured using *(i)* balanced sorbents that retain *(ii)* hydrophobic and hydrophilic compounds while *(iii)* preferentially removing phospholipids, salts and other unwanted matrix contaminants that cause column blockage; *(iv)* the workflow can be scaled-up using a 96 ‘blade’ cleanup system. **c**, After proteolytic digestion of the protein precipitate (organic crash), peptides can be subject to stable isotope labeling (e.g. TMT) and offline HPLC fractionation and/or phosphopeptide enrichment prior to nLC-MS using the same instrumentation.

Since the lower flow rates of nLC/MS necessitate longer gradients (∼30-45 min) compared with traditional metabolite microflow run times (∼15-30 min), real time positive/negative ion mode switching was applied during the MS/MS data acquisition to improve the efficiency and diversity of analyte detection. In parallel, the recovered total protein (organic precipitate) from the same samples were subjected to proteolysis (trypsin) and offline fractionation (basic reverse phase chromatography) prior to multiplex stable isotope labeling (Tandem Mass Tagging) prior to standard proteomic analysis using the same nLC/MS instrumentation (**Fig. 1c**).

The resulting paired spectral (MS1 and MS/MS) datasets are independently searched against protein and metabolite reference databases using stringent (*e.g.*1% false-discovery rate for protein identification) criteria to confidently identify biomolecules present in the respective cell compartments. The results of replicate and comparative analyses are subject to bioinformatic analysis to define technical variation as well as statistically-significant differences in relative abundance between distinct biological sample sets (**Fig. 1d**).

### PANAMA improves chromatographic separation compared with current methods

We evaluated the performance of nLC/MS for the reproducible chromatographic separation and characterization of complex cellular mixtures in comparison with that achieved using a standard analytical (micro-flow) mLC-MS procedure^15^. As a defined test material, we performed LLE on reference human plasma (SRM-1950, NIST), then subjected the metabolite fraction (supernatant) to replicate rounds of reverse phase chromatography using either a micro- flow column (2.1 mm i.d. x 100 mm packed with 1.8 μm pore size C18 resin) or nanoflow (75 μm i.d. x 25 cm with 1.8 μm C18) nanocolumn with or without prior sample cleanup. High throughput SPME auto-sampler was showed in **Fig. 2a**. Column back pressure was monitored throughout the entire LC/MS runtime.

**Figure 2.**
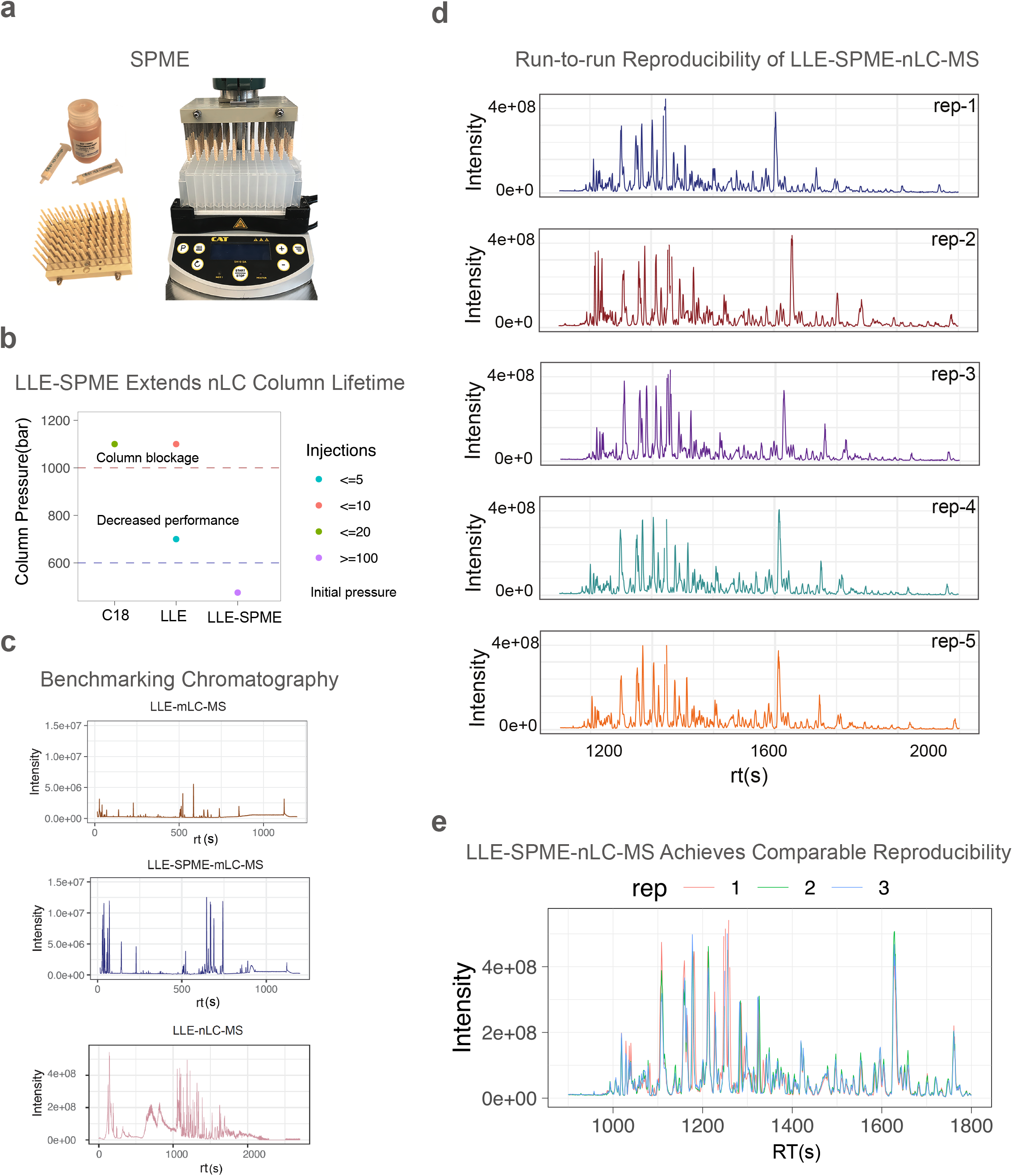
PANAMA benchmarking and performance evaluation **a,** Components of the multimodal open bed thin film SPME ‘blade’ capture system, including optimized sorbents (HLB SPE cartridge resin plus PS-DVB-WAX from CHROMABOND), coated blades and a 96-blade autosampler device (PAS Technology, Germany). **b,** Improved capillary column lifetime and reduced back pressure after SPME-based processing of a human plasma standard versus standard C18-tip cleanup or direct (LLE) injection. **c,** Side-by-side base peak chromatograms (BPC, positive ion mode) of human plasma standard extract with LLE and SPME by mLC-MS or nLC-MS. **d/e**, BPC traces of replicate nLC-MS injections after SPME pretreatment.

Notably, while LLE alone or C18 Sep-Pak sample processing resulted in nanocolumn blockage and rapid degradation performance, SPME pre-processing ensured stable nLC pressure over >100 sample injections (**Fig. 2b**). Regardless of flow rate, SPME significantly improved the chromatographic resolution as compared to the benchmarking micro-flow runs (**Fig. 2c&d**) and existing (low flow) approach ^10^. Despite the lower stability of nanospray ESI (**Fig. 2c&d**), SPME markedly improved the reproducibility of nLC-MS (**Fig. 2c/d/e**), resulting in lower coefficients of variation (% CV, largely below 15%) that are comparable with LLE alone (**Fig. 3a)**.

**Figure 3.**
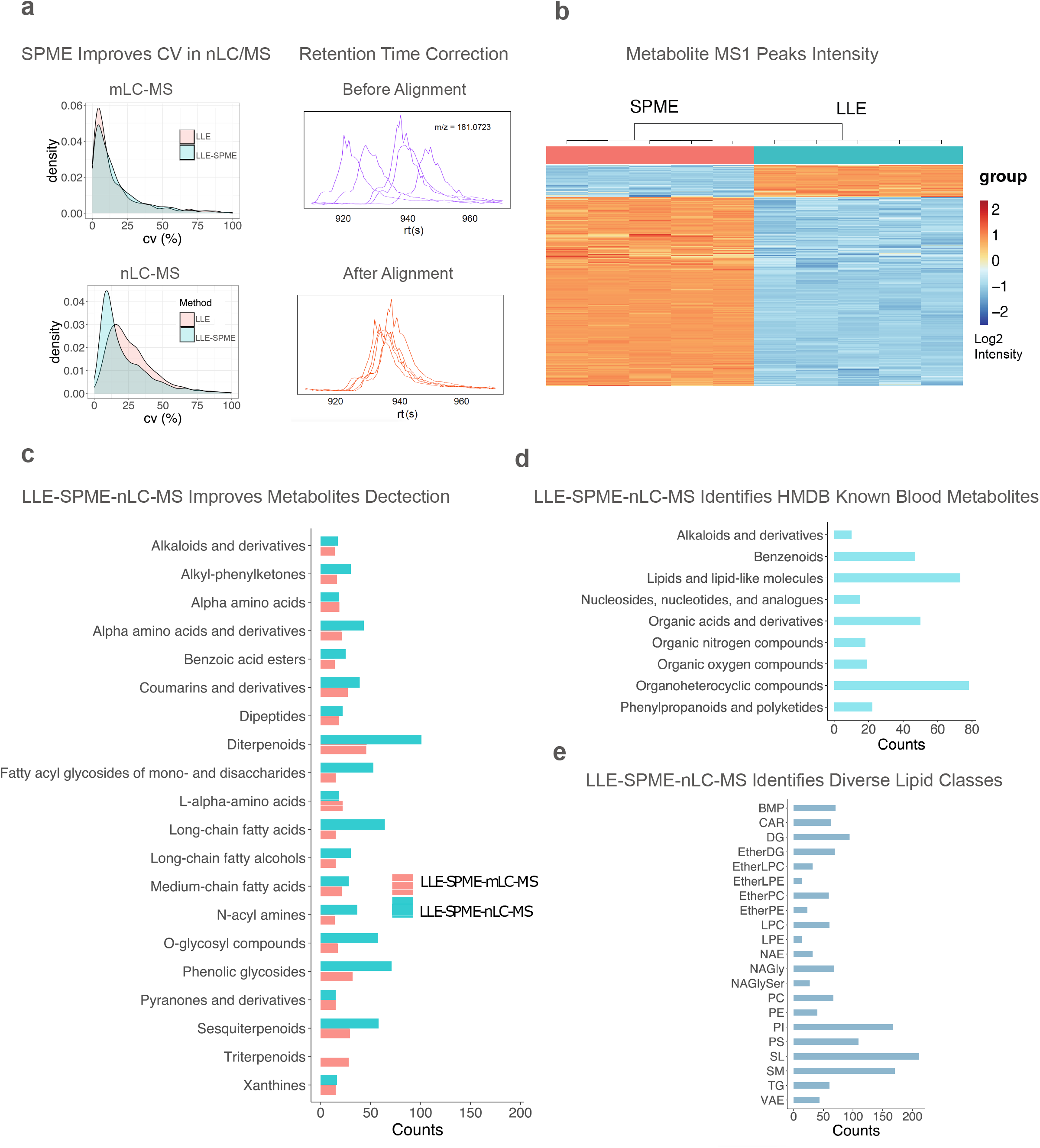
PANAMA coverage evaluation **a,** SPME cleanup reduced feature deviation (CV) in replicate plasma analyses by both nLC-MS and mLC-MS (pre- and post- alignment traces). **b,** Metabolite MS1 peak intensity regulation in LLE and SPME. **c**, Enriched metabolites (top 20 classes based on chemical similarity) detected by nLC-MS versus mLC-MS. PANAMA-based identification of known blood metabolites (**d**) and lipids (**e**).

Minor peak variability due to retention time drift was readily corrected by peak alignment (**Fig. 3a**). Regardless of flow rate, in comparison with LLE alone, SPME cleanup also markedly enhanced the signal recorded from a broad array of analytes (**Fig. 3b**). Strikingly, even half (49%) of the MS1 peaks detected by mLC-MS in positive ion mode showed at least a 2-fold increase in intensity after sample enrichment with negligible loss in total metabolite coverage (**Fig. 3c**, also see **Supplementary Table S1**).

### PANAMA achieves high molecular coverage

After assuring the robustness of the procedure, we next evaluated the effectiveness of SPME on global metabolite coverage achieved by nLC-MS as compared to current microflow. As summarized in **Supplementary Table S1**, high probability metabolites were identified using four community assessment criteria^16^.

Strikingly, nLC-MS with SPME sample processing detected nearly twice as many compounds (4026 *vs* 2084 level 2-3 metabolites) as compared to mLC-MS **(Supplementary Table S1)**. Moreover, while both flow rates produced similar compound profiles overall (**Fig. 3c**), the LLE-SPME-nLC-MS workflow generated higher total counts for 7 of the 20 top over- represented metabolite classes relative to only a slight loss of highly polar compounds (*eg.* amino acids). In total, PANAMA identified 448 annotated blood metabolites (Human Metabolome Database V4.0), including carboxylic acids, carbohydrates, nucleotides and vitamins (**Fig. 3d**).

Despite their high structural diversity, the LLE-SPME-nLC-MS workflow also identified 987 lipids using LipidBlast^17^ (**Fig. 3e**), including saccharolipids (SL), sphingolipids (*eg.* SM), glycerophospholipids (*eg.* LPC), and even low abundant glycerophosphoinositols (PI).

To improve the identification confidence, database matching was performed using MS- DIAL^18^ and a previously described^19^ decoy adduct search strategy for rigorous FDR estimation (**Fig. 4a**). As a further test of system performance, we spiked in 35 synthetic drug-like compounds into a reference blood specimen over concentration range of 4.3 pM to 1.67 μM (**Fig. 4b**). Whereas no search hits were detected in control (unspiked) plasma, all 35 exogenous compounds were readily identified in the spiked samples by PANAMA (based on high-energy collision dissociation spectral matching against an in-house compound reference library).

**Figure 4.**
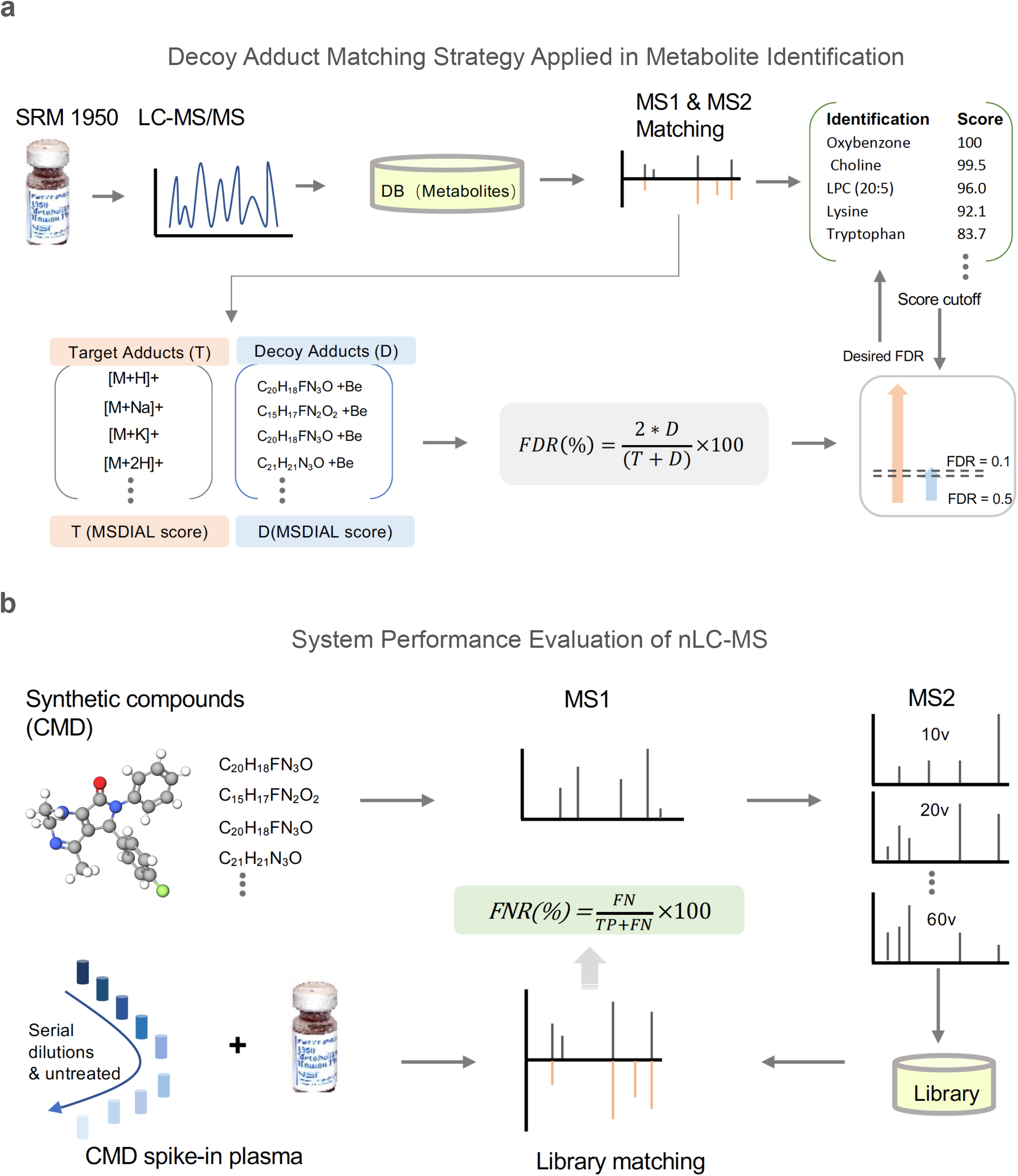
Schematics related to the estimation of the false negative and false positive rates **a,** Target and decoy adduct sets were applied using MS-DIAL annotation, and an empirically based score was then used as the cutoff for the desired FDR. **b,** Serial dilutions of synthetic compounds spiked into human plasma and identified using an in-house CMD library for FNR.

Absolute quantification analysis established an on-column limit-of-detection (LOD) of 2 to 5 fg with a linear precursor ion signal intensity over a large (x10^6^) dynamic range (**Supplementary Table S1**).

### Application of PANAMA to complex cell lysates

To further evaluate the utility of our workflow in different pathobiological contexts, we used SPME-nLC/MS to examine the global proteomic and metabolomic changes that occur in breast cancer cells (MB-MDA-231) in response to acute glucose withdrawal in culture, which leads to a well characterized adaptive response^20^. Analysis of the resulting multi-omic profiling data using principal component analysis (PCA) revealed reproducible but distinct metabolite and protein-based cellular states (profile groupings) in response to low glucose (**Fig. 5a**).

**Figure 5.**
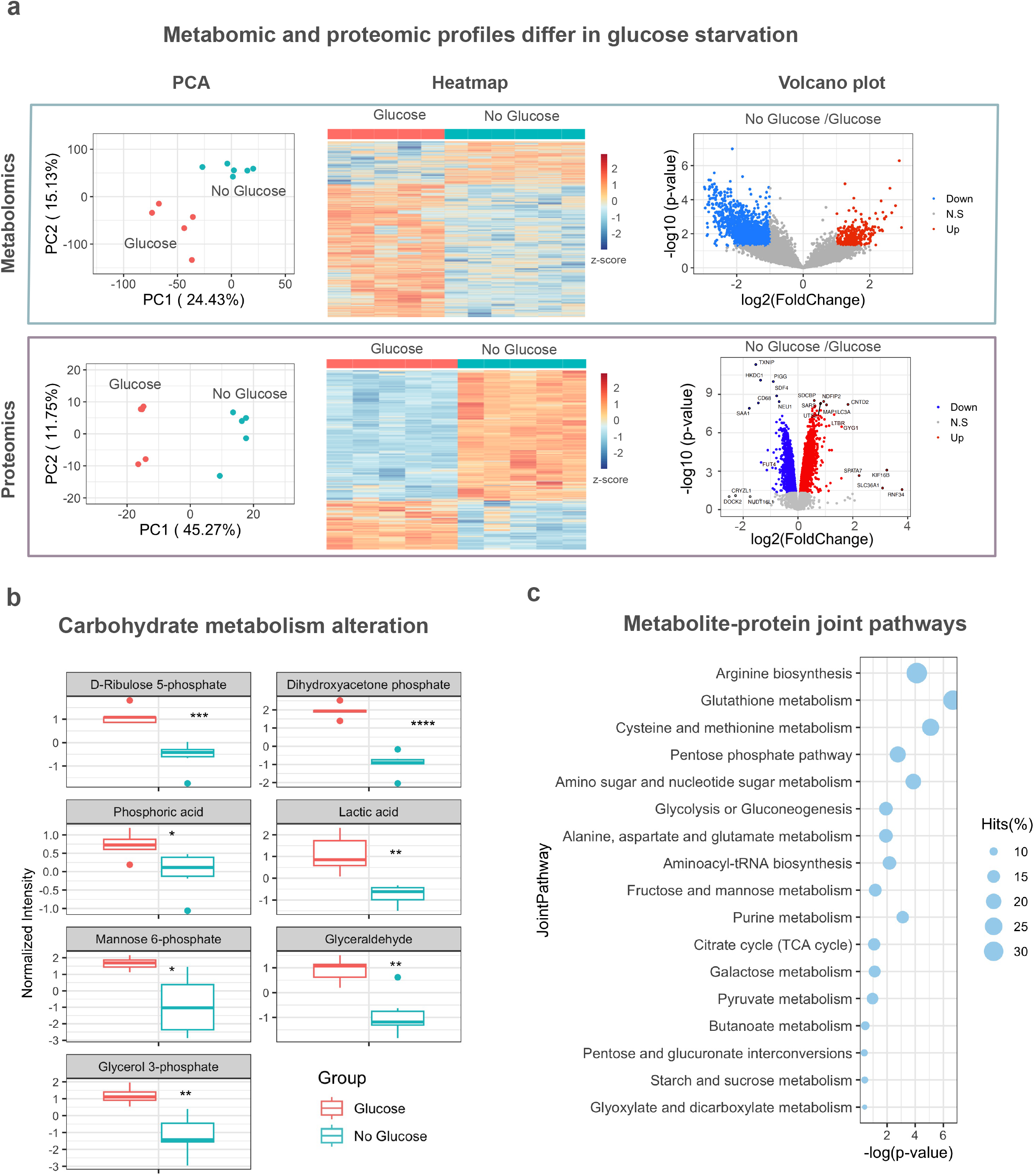
Application of PANAMA to study remodeling of cellular metabolism **a**, PCA plot, heatmap and volcano plots of PANAMA-based metabolomic and proteomic profiles of breast cancer cells identified under glucose starvation versus standard culture conditions. **b**, Box plots of differential metabolites in cancer cells associated with carbohydrate metabolism in response to glucose starvation. **c**, Joint metabo-proteomic enriched (KEGG) pathways based on parallel analyses.

Notably, our streamlined approach for jointly measuring metabolomic and proteomic profiles identified a comparable number of proteins as compared with previous surveys of the same cell line^21^. As negative breast cancer especially depends on glucose metabolism^22^, we identified dozens of significantly upregulated (p<0.05, FC>1.5) proteins associated with amino sugar metabolic process, glucose catabolic process to pyruvate, glycolytic process through fructose-6-phosphate and carbohydrate biosynthetic process (**Supplementary Table S2)**, which had roles in metabolic compensation in response to starvation. For instance, uridine phosphorylase 1 (UPP1), enzyme that catalyzes carbon and energy sources responding to glucose starvation^23^ and glycogenin-1 (GLYG, see volcano plot in **Fig .5a**), enzyme that involves in glucosyltransferase activity^24^.

Although cancer cell produces energy through glycolysis (Warburg effect) rather than mitochondrial oxidative phosphorylation, acute glucose starvation (24h) may induce oxidative stress response^25^. Surprisingly, we identified many proteins related to mitochondrial translational elongation, transmembrane transport and mitochondrial gene expression were significantly suppressed (FDR<0.05, see **Supplementary Table S2**), revealing glucose starvation (48h) sensitized the breast cancer cell to death/apotosis via mitochondrial dysfunction. This effect also indicated by the upregulation of many proteins that lead to apoptotic process, including E3 ubiquitin-protein ligase (RNF34), caspase recruitment domain-containing protein 6 (CARD6), and tumor necrosis factor receptor superfamily member 3 (LTBR) (see volcano plot in **Fig. 5a**)

Contrarily, 400 of the 3446 annotated (level 3) metabolites identified using PANAMA (**Supplementary Table S2**) were found to be significantly downregulated (*p*<0.05, ≥1.5-fold- change) upon glucose withdrawal as compared to growth in standard glucose-containing media (**Fig. 5a**). Once again, many of these differential molecules, such as d-ribulose 5-phosphate, dihydroxyacetone phosphate and phosphoric acid, lactic acid, mannose 6-phosphate, glyceraldehyde and glycerol 3-phosphate **(Fig. 5b),** were associated with fructose and mannose metabolism/degradation, Warburg effect, gluconeogenesis and glycolysis, even mitochondrial electron transport chain.

To gain a more comprehensive picture of the molecular change response to glucose starvation, we performed metabo-proteomic joint-pathway enrichment using altered markers. Our integrated analysis revealed many molecules related to carbohydrate metabolism, including pentose phosphate pathway, amino sugar and nucleotide sugar metabolism and glycolysis or gluconeogenesis, galactose metabolism, pyruvate and butanoate metabolism **(Fig. 5c)** along with TCA cycle and compounds linked to arginine biosynthesis, which associated with cell proliferation^26^, glutathione metabolism and cysteine and methionine metabolism, the non- essential amino acids that become essential for tumor proliferation and survival^27^**(Fig. 5c)**.

These results show that our unified nanoflow-based sample preparation workflow can provide complementary insights into global biochemical rewiring of cell metabolism.

## Discussion

PANAMA represents an optimized, practical and robust sampling technique that enables metabolomic (lipidomic) analysis using nanoflow procedures that empower virtually all current proteomics approaches. By eliminating interfering matrix components, particularly phospholipids, that cause ion suppression and poor spray stability, SPME-nLC/MS is well suited to broad array of ‘dirty’ samples ranging from whole cell lysates, biofluids such as blood, and tissue extracts.

While SPME has previously been used to process serum samples prior to analytical flow (mLC- MS) protocols^13^, to our knowledge its combination with true nanoflow ESI has not been reported beyond our group. We established that SPME dramatically extends capillary column life times, while still enabling detection of diverse compound types for untargeted metabolomics.

As shown in independent benchmarking and use case studies, our unified ‘one-pot’ PANAMA workflow measures metabolites, lipids and proteins with comparable quality and depth as for more traditional analytical workflows optimized for different molecular classes. While not intended as a *replacement* for established analytical chemistry workflows, we note that PANAMA achieves notably high sensitivity and chromatographic resolution with acceptable inter-assay CVs, all while empowering deep proteomic analysis in parallel. We illustrated this complementarity by examining metabolic pathway remodeling at the level of both enzymes and small molecules in a cancer cell model.

## Limitations of the approach

Our unified methodology provides complementary biomolecular information (metabolite- lipid-protein) for biospecimens using the same exact instrumentation, bypassing the new for additional sample consumption or redundant infrastructure. A potential drawback is loss of certain analytes (*eg.* high polar amino acids) during sample cleanup. Moreover, as is commonly observed in the field, it remains challenging to capture the full chemical diversity of cellular metabolites in a single chromatographic separation ^28^. But we deem these as acceptable trade- offs given the enhanced robustness and overall high discovery potential exhibited by PANAMA.

While global (untargeted) data acquisition is a preferred mode of operation, the SPME- nLC workflow described here can be tailored for target directed or data-independent analyte detection by incorporation of diagnostic fragment ions and spectral library matching. This flexibility is partially offset by the lack of commercially available metabolite reference standards and annotated MS/MS spectra, which hinder confident isomer discrimination^29^.

## STAR Methods

### Chemical and materials

LC-MS grade solvent were obtained from Fisher Scientific. Frozen human plasma standard (SRM1950) was purchased from NIST (USA). Polystyrene-divinylbenzene (PS-DVB) and N-vinyl pyrrolidone were purchased from Macherey Nagel (USA). Weak anion exchange particles were obtained from HLB SPE cartridges (Waters, USA). The SPME blade Unit and robotic 96 autosampler were purchased from Professional Analytical Systems (PAS) Technology (Magdala, Germany) for metabolomic crude extracts clean-up.

### Cancer cell line culture

The human breast cancer-derived cell line MDA-MB-231 was obtained from the American Type Culture Collection (ATCC). The cells were maintained in a humidified incubator at 37°C with 5% CO_2_ in Dulbecco’s modified Eagle’s medium (DMEM) without pyruvate containing 4.5 g/L glucose and L-glutamine (Gibco) supplemented with 10% heat inactivated fetal bovine serum (FBS; HyClone) and 100 units/mL penicillin-100 µg/mL streptomycin (HyClone). Cells were then either continued to be cultured in the same medium or were switched to a no glucose condition, which was DMEM containing L-glutamine but without glucose or pyruvate for 48 hours before harvesting.

### Metabolite extraction

Human plasma standard (100μL, SRM1950, NIST), or MDA-MB-231 cell (12 million) pellets were resuspended in a 4 vol. mixture of ice-cold methanol/acetonitrile/water (MeOH/ACN/H_2_O, 40/40/20 v/v; MS-grade, Fisher) in chemically resistant microcentrifuge tubes (*eg.* Eppendorf) and vortexed for 30 s. The samples were flash frozen in liquid nitrogen for 1 min, allowed to thaw and disrupted on ice by a probe sonicator (40 kHz, Ultra Autosonic) at 10% power for 5 min; this cycle was performed three times (note that sonication was not performed for biofluid extracts). Afterward, the extracts were incubated for 1h at −20°C and centrifuged at 12,000 x g and 4°C for 15 min to pellet the protein precipitate. The metabolite-containing supernatants were carefully transferred to new microtubes, dried under vacuum at 30°C, and kept at -80°C prior to LC-MS. For LC-MS analysis, extracts were thawed and resolubilized in 200 μL of 2% methanol and subjected to SPME (see next section) for matrix cleanup. The protein precipitate was maintained at 4°C prior to tryptic digestion and proteomic analysis. Equal amounts of each sample were pooled as an internal quality control (QC).

### Solid-phase microextraction (SPME)

The metabolite mixtures were transferred to a 96-well microwell plate for SPME processing (**Supplementary Fig. S1a**). The coated blades were washed with EtOH/H_2_O (70:30, v/v) for 30 min and preconditioned for 30 min in MeOH/H_2_O (50:50, v/v). The samples were extracted by incubation with the blades for 1 h at room temperature. The blades were briefly rinsed for ∼20 s using water, and then the bound metabolites were desorbed using ACN/H_2_O (50:50, v/v) for 1 h. After the solvent was evaporated to dryness using a vacuum concentrator at 30°C, the samples were reconstituted in 2% ACN for analysis by LC-MS. For the performance evaluation, 5-folds serial dilutions of mixture of 35 synthetic compounds (Boston University Center for Molecular Discovery) from a concentration of 1.67 μM were spiked into SPME-cleaned up human plasma (SRM1950) extracts. **Supplementary Table S1** provides a list of the synthetic compounds along with their masses and structures.

### LC-MS analysis of metabolites

Metabolite nanoflow (nLC) MS analysis was performed using a Hybrid Quadrupole- Orbitrap Q-Exactive HF (Thermo Scientific) with C18 reversed-phase pre-column (75 mm i.d. × 2 cm, 3μm) and capillary column (75 mm i.d. × 2 cm, 2μm, 100 Å, ThermoFisher Scientific); the column oven was set to 40°C. To prevent ion suppression in the negative mode, formic acid was not used in the mobile phase; trifluoroacetic acid (TFA, a common contaminant from phosphopeptide enrichment) was similarly avoided. The mobile phase A was 2% ACN, and mobile phase B was 80% ACN. The nLC flow rate was 300 nL/min. Samples (2μL) were injected and separated over a 45 min gradient. The gradient consisted of 2% to 60% mobile phase B for 20 min, was increased to 95% mobile phase B over 10 min and maintained at 95% mobile phase B for 15 min. The MS instrument was operated in the automated switching ESI mode over a full mass scan range of m/z 67−1000 at a resolution of 60,000. The AGC target was set to 3×e^6^ ions, and the maximum ion injection time was set to 25 ms. The source ionization parameters were optimized for a transfer temperature at 300°C and a spray voltage set to 2.1 kV and -1.8 kV for the positive and negative modes, respectively. MS2 scans were performed at 15,000 resolution with a maximum injection time of 64 ms using stepped normalized collision energies (NCEs) of 10, 20, and 40. Dynamic exclusion was enabled using a time window of 10 s.

A parallel analysis by micro-flow LC-MS was performed using a 6456 Q-TOF mass spectrometer coupled to a 1290 Infinity II LC system (Agilent Technologies). Plasma aliquot with/without SPME cleaned up (2μL) were loaded and seperated on an Agilent Zorbax C-18 UHPLC column (100 x 2.1 mm; 1.8 μm) over a 20-min gradient. Following an MS1 full scan with a spectrometry range of 50-1100 m/z in the positive mode, the precursors were fragmented using multiple collision energies of 10, 20 and 40.

### Protein digestion

After pelleting, the protein precipitates from the organic solvent extraction^8^ were resuspended in 250 μL of lysis buffer containing 6 M guanidine hydrochloride (GuHCl), protease inhibitors (Sigma) and phosphatase inhibitors (Roche). The samples were heated at 95°C for 10 min, cooled on ice for 10 min and then briefly sonicated to shear the nucleic acids. The samples were diluted with 100 mM Tris (pH 8.5) to reduce the concentration of GuHCl to 0.75 M. After quantification with a BCA kit (Thermo Scientific), the proteins were digested overnight with sequence-grade trypsin (enzyme-to-protein ratio of 1:50) at 37°C, and formic acid was then added to obtain a final concentration of 1% in solution. The resulting peptides were desalted using a C18 Sep-Pak (Waters) according to the manufacturer’s instructions.

### TMT peptide labeling

Prior to tandem mass tag (TMT) labeling, peptide quantification was performed by Pierce quantitative colorimetric assay (Thermo Scientific). According to the manufacturer’s instructions, 100 μg of peptide per sample was resuspended in 0.1 M triethylammonium bicarbonate (TEAB). Peptides of cancer cell (5 channels per condition, glucose & no glucose) were labeled with 0.2 mg of TMT 10-plex while 15 samples of human serum (5 channels per condition, healthy, pre- and post-dialysis) were labeled with TMTpro (Thermo Scientific) for 1 h at room temperature. To quench the reaction, 5% hydroxylamine was added to each sample, and the resulting mixture was incubated at room temperature for 15 min. After labeling, equal amounts of each sample were combined in a new microtube and desalted using a C18 Sep-Pak (Waters).

### High-pH reversed-phase peptide fractionation

Peptides (500 μg) were fractionated offline on a Waters XBridge BEH C18 reversed- phase column (3.5 μm, 4.6 ×250 mm) using an Agilent 1100 HPLC system operated at a flow rate of 0.45 mL/min with two buffer lines: buffer A (consisting of 0.1% ammonium hydroxide-2% acetonitrile-water) and buffer B (consisting of 0.1% ammonium hydroxide-98% acetonitrile, pH 9). The peptides were separated by a gradient from 0% to 10% B in 5 min followed by linear increases to 30% B in 23 min, to 60% B in 7 min, and then 100% in 8 min and maintained at 100% for 5 min. This separation yielded 48 collected fractions that were subsequently combined into 12 fractions and then evaporated to dryness in a vacuum concentrator. The peptides (2 µg) from each fraction were reconstituted in 0.1% formic acid and maintained at -80°C prior to analysis by nLC-MS/MS.

### LC-MS analysis of peptides

LC-MS analysis was performed using same Q-Exactive HF system as described above.

Peptides were loaded onto a C18 pre-column (75 mm i.d. × 2 cm, 100 Å, ThermoFisher Scientific) then separated on a reverse-phase nano-spray column (75 mm i.d. × 50 cm, 100 Å, ThermoFisher Scientific) using gradient elution. Peptdies (2 μg) were injected and separated over 150 min gradient. The mobile phase A was consisted of 0.1% FA-2% ACN-Water, and mobile phase B was consisted of 0.1% FA-80% ACN-Water. The gradient consisted of 6% to 40% mobile phase B over 155 min, was increased to 95% mobile phase B over 4 min and maintained at 95% mobile phase B for 3 min at a flow rate of 250 nL/min. The MS instrument was operated in positive ion mode over a full mass scan range of m/z 350−1400 at a resolution of 60,000 with a normalized AGC target of 300%. The source ion transfer tube temperature was set at 275°C and a spray voltage set to 2.5kv. Data was acquired on a data dependent mode with FAIMS running 3 compensation voltages at -50v, -57v and -64v. MS2 scans were performed at 45,000 resolution with a normalized collision energy 34. Dynamic exclusion was enabled using a time window of 60 s.

### Metabolomic data processing

The raw chromatographic data files were converted to mzML format and split into positive and negative ion mode files with msConvert^30^ prior to analysis using MS-DIAL (V4.18). ‘Linear-weighted moving average’ was used for peak detection, and the minimum peak height was set to 2000. Afterward, spectral centroiding was performed by integrating the mass spectrum over the ±0.01 and ±0.025 Da range in MS1 and MS2, respectively. Common adducts (*e.g.*, [M+H]^+^, [M+NH_4_]^+^, [M+Na]^+^, [M-H]^-^, [M-H_2_O-H]^-^, and [M+Cl]^-^) were annotated prior to identification. Implausible adducts (*e.g.*, [M+Be]^+/-^) were defined as decoy targets for FDR estimation and searched along with the native adducts to define a suitable matching score cutoff (*e.g.*, >0.7) to improve metabolite identification reliability (*e.g.*, decoy-over-native adduct ratio <0.1). The spectra were searched using MS-DIAL^18^ against a metabolomic (MSMS-Public-Pos-VS15.msp) or lipidomic (LipidMsmsBinaryDB-VS68-FiehnO.lbm2) library with a matched mass tolerance of 0.025 and 0.05 Da for MS1 and MS2 ions, respectively. The QC samples were specified as reference files for sample alignment. The retention time shift was evaluated on the basis of the extracted ion chromatogram (EIC) of single identified compounds prior to assigning an RT tolerance window. The analysis of paraxanthine (m/z 181.0723, delta ppm=1.7) is shown in **Supplementary Fig. S1g&h** as an example to illustrate the process. The data matrixes were exported as tab-delimited text files. Features with high CV intensity (≥50), low fold change (≤5) in average sample intensity relative to negative controls, low signal/noise (≤3), and decoy adducts ([M+Be]^+^ and [M+Be]^2+^) were removed prior to subsequent downstream analysis. MetaboAnalyst 5.0^31^ with standard parameters was used for pathway enrichment.

### Proteomic data analysis

MS2 spectra were processed and searched by MaxQuant (version 1.6) against a database containing native (forward) human protein sequences (UniProt) and reversed (decoy) sequences for protein identification. The search allowed for two missed trypsin cleavage sites, variable modifications of methionine oxidation, and N-terminal acetylation. The carbamidomethylation of cysteine residues was set as a fixed modification. Ion tolerances of 20 and 6 ppm were set for the first and second searches, respectively. The candidate peptide identifications were filtered assuming a 1% FDR threshold based on searching the reverse sequence database. Quantification was performed using the TMT reporter on MS2 (TMT10- plex). Bioinformatic analysis was performed in the R statistical computing environment (version 3.6.1). Enrichment analysis was performed using the web-based tool MetaboAnalyst 5.0 against the KEGG database (https://www.genome.jp/kegg/).

## Supporting information

Supplementary Protocol 1

## Acknowledgements

The authors are grateful for the assistance and critical input provided by colleagues in Toronto and Boston. This work was supported by operating funds to A.E. from the Canadian Institutes of Health Research (FDN-148399) and generous start-up funds from Boston University. The authors thank John A. Porco, Lauren E. Brown, and Han Yueh for providing synthetic compounds and Kieran Wynne for providing technical support on mass spectrometry. The authors thank Pietro Morlacchi, Alex Hansler, and Olivia Cohen from Agilent Technologies Inc. for the technical assistance provided. The authors thank Juntuo Zhou from Peking University for the technical suggestion on metabolomics.

## Author contributions

A.E. conceived and supervised the study. W.L. and F.M. developed and optimized the metabolite extraction protocol and the nLC-MS method and parameters. W.L. performed the data acquisition. W.L., B.C.B. and C.F.H. performed the data analyses. N.L. contributed to sample preparation, R.H., H.G. and M.M provided technical support. W.L., B.C.B., and A.E. cowrote the manuscript.

## Data availability

Data for this project have been deposited to the MassIVE archive accession code MSV000091579 (doi:10.25345/C5GX4548D).

## Supplemental data

This article contains supplemental data

## References

1. Dubin, R.F., and Rhee, E.P. (2020). Proteomics and Metabolomics in Kidney Disease, including Insights into Etiology, Treatment, and Prevention. Clin J Am Soc Nephrol 15, 404–411. 10.2215/CJN.07420619.

2. Merbel, N.C.v.d. (2019). Protein quantification by LC-MS: a decade of progress through the pages of Bioanalysis. Bioanalysis 11, 629–644. 10.4155/bio-2019-0032.

3. Gika, H., Virgiliou, C., Theodoridis, G., Plumb, R.S., and Wilson, I.D. (2019). Untargeted LC/MS-based metabolic phenotyping (metabonomics/metabolomics): The state of the art. J Chromatogr B Analyt Technol Biomed Life Sci 1117, 136–147. 10.1016/j.jchromb.2019.04.009.

4. Dupree, E.J., Jayathirtha, M., Yorkey, H., Mihasan, M., Petre, B.A.-O., and Darie, C.A.- O. A Critical Review of Bottom-Up Proteomics: The Good, the Bad, and the Future of this Field. Proteomes 8, 2227–7382. 10.3390/proteomes8030014.

5. Han, X., and Gross, R.W. (2005). Shotgun lipidomics: electrospray ionization mass spectrometric analysis and quantitation of cellular lipidomes directly from crude extracts of biological samples. Mass Spectrom Rev 24, 367–412. 10.1002/mas.20023.

6. Kofeler, H.C., Ahrends, R., Baker, E.S., Ekroos, K., Han, X., Hoffmann, N., Holcapek, M., Wenk, M.R., and Liebisch, G. (2021). Recommendations for good practice in MS- based lipidomics. J Lipid Res 62, 100138. 10.1016/j.jlr.2021.100138.

7. Fischer, R., Bowness, P., and Kessler, B.M. (2013). Two birds with one stone: doing metabolomics with your proteomics kit. Proteomics 13, 3371–3386. 10.1002/pmic.201300192.

8. Causon, T.J., and Hann, S. (2016). Review of sample preparation strategies for MS-based metabolomic studies in industrial biotechnology. Anal Chim Acta 938, 18–32. 10.1016/j.aca.2016.07.033.

9. Telu, K.H., Yan, X., Wallace, W.E., Stein, S.E., and Simon-Manso, Y. (2016). Analysis of human plasma metabolites across different liquid chromatography/mass spectrometry platforms: Cross-platform transferable chemical signatures. Rapid Commun Mass Spectrom 30, 581–593. 10.1002/rcm.7475.

10. Simon-Manso, Y., Lowenthal, M.S., Kilpatrick, L.E., Sampson, M.L., Telu, K.H., Rudnick, P.A., Mallard, W.G., Bearden, D.W., Schock, T.B., Tchekhovskoi, D.V., et al. (2013). Metabolite profiling of a NIST Standard Reference Material for human plasma (SRM 1950): GC-MS, LC-MS, NMR, and clinical laboratory analyses, libraries, and web- based resources. Anal Chem 85, 11725–11731. 10.1021/ac402503m.

11. Moore, J., Ewoldt, J., Venturini, G., Pereira, A.C., Padilha, K., Lawton, M., Lin, W., Goel, R., Luptak, I., Perissi, V., et al. (2023). Multi-Omics Profiling of Hypertrophic Cardiomyopathy Reveals Altered Mechanisms in Mitochondrial Dynamics and Excitation-Contraction Coupling. Int J Mol Sci 24. 10.3390/ijms24054724.

12. Deehan, M., Lin, W., Blum, B., Emili, A., and Frydman, H. (2021). Intracellular Density of Wolbachia Is Mediated by Host Autophagy and the Bacterial Cytoplasmic Incompatibility Gene cifB in a Cell Type-Dependent Manner in Drosophila melanogaster. mBio 12. 10.1128/mBio.02205-20.

13. Mousavi, F., Bojko, B., and Pawliszyn, J. (2015). Development of high throughput 96- blade solid phase microextraction-liquid chromatrography-mass spectrometry protocol for metabolomics. Anal Chim Acta 892, 95–104. 10.1016/j.aca.2015.08.016.

14. Mirnaghi, F.S., and Pawliszyn, J. (2012). Development of coatings for automated 96-blade solid phase microextraction-liquid chromatography-tandem mass spectrometry system, capable of extracting a wide polarity range of analytes from biological fluids. J Chromatogr A 1261, 91–98. 10.1016/j.chroma.2012.07.012.

15. Wang, L., Naser, F.J., Spalding, J.L., and Patti, G.J. (2019). A Protocol to Compare Methods for Untargeted Metabolomics. Methods Mol Biol 1862, 1–15. 10.1007/978-1-4939-8769-6_1.

16. Sumner, L.W., Amberg, A., Barrett, D., Beale, M.H., Beger, R., Daykin, C.A., Fan, T.W., Fiehn, O., Goodacre, R., Griffin, J.L., et al. (2007). Proposed minimum reporting standards for chemical analysis Chemical Analysis Working Group (CAWG) Metabolomics Standards Initiative (MSI). Metabolomics 3, 211–221. 10.1007/s11306-007-0082-2.

17. Kind, T., Liu, K.H., Lee, D.Y., DeFelice, B., Meissen, J.K., and Fiehn, O. (2013). LipidBlast in silico tandem mass spectrometry database for lipid identification. Nat Methods 10, 755–758. 10.1038/nmeth.2551.

18. Tsugawa, H., Cajka, T., Kind, T., Ma, Y., Higgins, B., Ikeda, K., Kanazawa, M., VanderGheynst, J., Fiehn, O., and Arita, M. (2015). MS-DIAL: data-independent MS/MS deconvolution for comprehensive metabolome analysis. Nat Methods 12, 523–526. 10.1038/nmeth.3393.

19. Palmer, A., Phapale, P., Chernyavsky, I., Lavigne, R., Fay, D., Tarasov, A., Kovalev, V., Fuchser, J., Nikolenko, S., Pineau, C., et al. (2017). FDR-controlled metabolite annotation for high-resolution imaging mass spectrometry. Nat Methods 14, 57–60. 10.1038/nmeth.4072.

20. Graham, N.A., Tahmasian, M., Kohli, B., Komisopoulou, E., Zhu, M., Vivanco, I., Teitell, M.A., Wu, H., Ribas, A., Lo, R.S., et al. (2012). Glucose deprivation activates a metabolic and signaling amplification loop leading to cell death. Mol Syst Biol 8, 589. 10.1038/msb.2012.20.

21. Alghanem, B., Ali, R., Nehdi, A., Al Zahrani, H., Altolayyan, A., Shaibah, H., Baz, O., Alhallaj, A., Moresco, J.J., Diedrich, J.K., et al. (2020). Proteomics Profiling of KAIMRC1 in Comparison to MDA-MB231 and MCF-7. Int J Mol Sci 21, 4328. 10.3390/ijms21124328.

22. Shin, E., and Koo, J.S. (2021). Glucose Metabolism and Glucose Transporters in Breast Cancer. Front Cell Dev Biol 9, 728759. 10.3389/fcell.2021.728759.

23. Watanabe, S., and Uchida, T. (1995). Cloning and expression of human uridine phosphorylase. Biochem Biophys Res Commun 216, 265–272. 10.1006/bbrc.1995.2619.

24. Chaikuad, A., Froese, D.S., Berridge, G., von Delft, F., Oppermann, U., and Yue, W.W. (2011). Conformational plasticity of glycogenin and its maltosaccharide substrate during glycogen biogenesis. Proc Natl Acad Sci U S A 108, 21028–21033. 10.1073/pnas.1113921108.

25. Raut, G.K., Chakrabarti, M., Pamarthy, D., and Bhadra, M.P. (2019). Glucose starvation-induced oxidative stress causes mitochondrial dysfunction and apoptosis via Prohibitin 1 upregulation in human breast cancer cells. Free Radic Biol Med 145, 428–441. 10.1016/j.freeradbiomed.2019.09.020.

26. Chen, C.L., Hsu, S.C., Ann, D.K., Yen, Y., and Kung, H.J. (2021). Arginine Signaling and Cancer Metabolism. Cancers (Basel) 13. 10.3390/cancers13143541.

27. Combs, J.A., and DeNicola, G.M. (2019). The Non-Essential Amino Acid Cysteine Becomes Essential for Tumor Proliferation and Survival. Cancers (Basel) 11. 10.3390/cancers11050678.

28. I. Barnes, S., Benton, H.P., Casazza, K., Cooper, S.J., Cui, X., Du, X., Engler, J., Kabarowski, J.H., Li, S., Pathmasiri, W., et al. (2016). Training in metabolomics research. Designing the experiment, collecting and extracting samples and generating metabolomics data. J Mass Spectrom 51, 461–475. 10.1002/jms.3782.

28. Chaleckis, R., Meister, I., Zhang, P., and Wheelock, C.E. (2019). Challenges, progress and promises of metabolite annotation for LC-MS-based metabolomics. Curr Opin Biotechnol 55, 44–50. 10.1016/j.copbio.2018.07.010.

29. Adusumilli, R., and Mallick, P. (2017). Data Conversion with ProteoWizard msConvert. Methods Mol Biol 1550, 339–368. 10.1007/978-1-4939-6747-6_23.

30. Chong, J., and Xia, J. (2020). Using MetaboAnalyst 4.0 for Metabolomics Data Analysis, Interpretation, and Integration with Other Omics Data. Methods Mol Biol 2104, 337–360. 10.1007/978-1-0716-0239-3_17.

