## Supplementary Protocol 1 for "PANAMA enabled high sensitivity dual nanoflow LC/MS metabolomics and proteomics analysis"

**Solid phase microextraction (SPME) coating preparation procedure**

**Blades pretreatment**

Prior to the coating process, the blades were sonicated with concentrated hydrochloric acid for 60 min to condition the stainless steel surface for effective immobilization of the coating (Figure p1). The blades were then washed with nanopure water. Next, Blades were covered with aluminum foil, and dried in an oven for 30 min at 150 ˚C and then cooled to room temperature. Upon coating preparation, 2 cm end of the blades were measured as Figure p2 and the other parts of the blades were covered by aluminum foil (Figure p2).

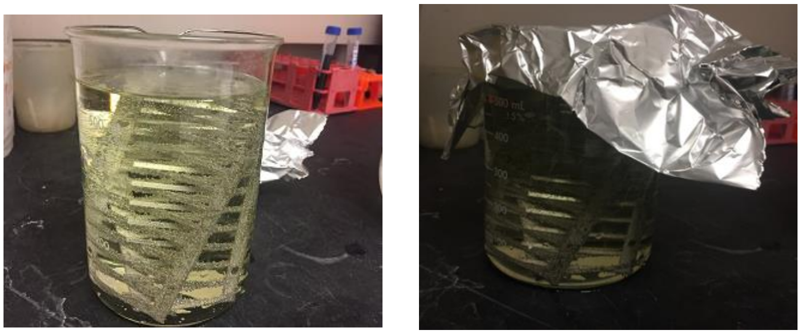

**Figure p1.** Blades precondition with hydrochloric acid for 60 min.

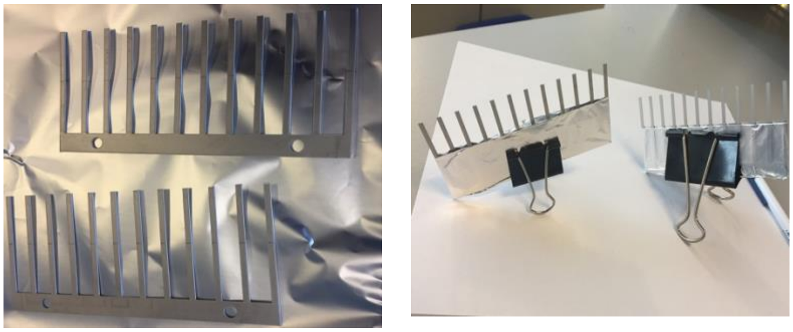

**Figure p2.** Preparation of blades (2cm end) for coating

**Preparation of biocompatible polyacrylonitrile (PAN) glue**

10% w/w PAN particles with N,N-dimethylformamide (DMF) solvent resulted in the optimum properties of the required glue. The mixture was heated in the oven at 90 ˚C for 1 h to dissolve, until a yellowish clear solution was obtained (Figure p3), then cooled to room temperature.

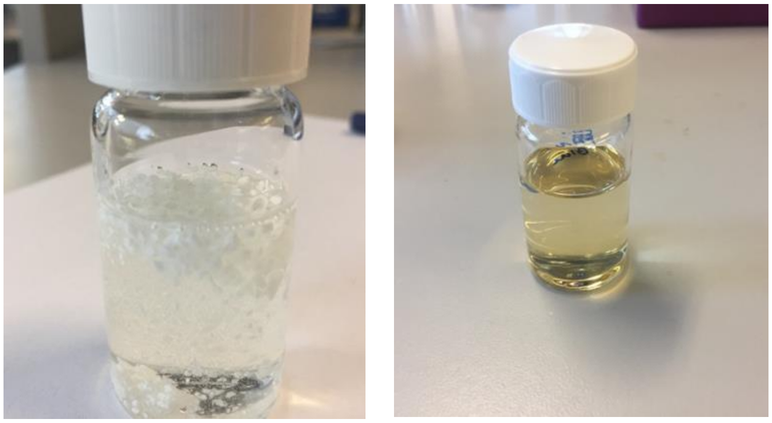

**Figure p3.** PAN in DMF (before and after) dissolve in solvent

### Coating particles on the stainless steel blades

Mix particles polystyrene-divinylbenzene with a weak anion exchanger (PS-DVB-WAX) and N-vinyl pyrrolidone (DVB-NVP) with 1:1 (w/w). Slurry of particles in prepared PAN glue was transferred into a glass sprayer flask. The source of nitrogen gas with the required pressure was connected to the sprayed and blades were coated by immobilizing particles on the surface of 96-blades. Particles, glass type sprayer, and sprayer set up connection to the nitrogen gas tank are demonstrated in Figure p4.

The coating preparation was performed by spraying very thin layers of particles slurry on the first 2 cm length of the blades followed by instant thermal curing in the oven at 180˚C for 2 min. The coating and curing steps were repeated 10 times in order to ensure uniform coverage and proper thickness of the coating on the surface of the blades (Figure p5, Video 1).

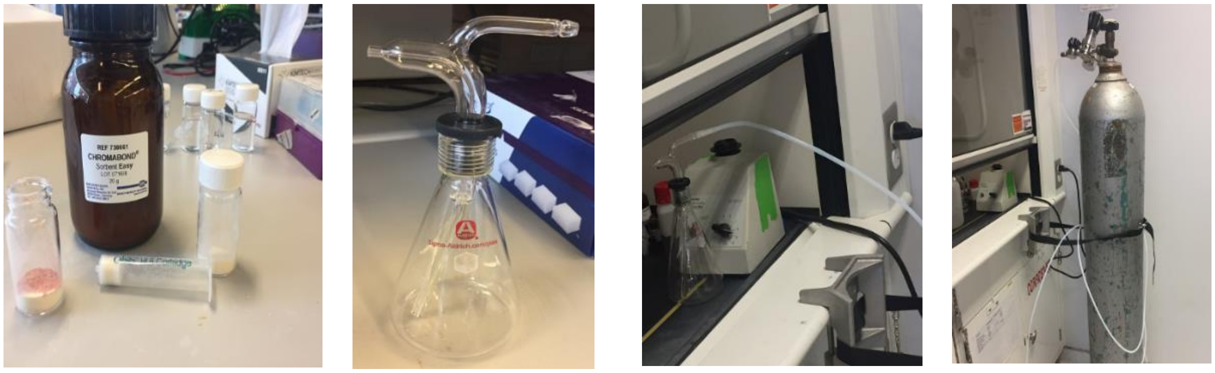

**Figure p4.** Bulk of PS-DVB-WAX from CHROMABOND and HLB SPE cartridges; Sigma flask type sprayer.

**Note) Coating procedure must be done under fume hood because of the chemical safety.

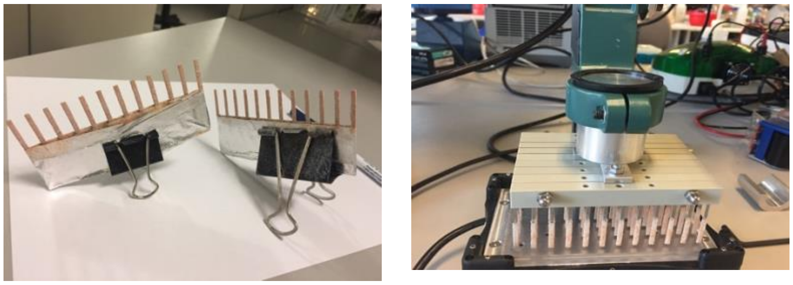

**Figure p5.** Coated blades; 96-concept autosampler
